# Region-specific modulation and predictive potential of the oscillatory dynamics in an *ex vivo* model of ictogenesis in the CA1, CA3 and dentate gyrus

**DOI:** 10.1101/2024.08.23.609340

**Authors:** Lida-Evmorfia Vagiaki, Dionysios Xydias, Maria Kefalogianni, Sotiris Psilodimitrakopoulos, Emmanuel Stratakis, Kyriaki Sidiropoulou

## Abstract

The hippocampus, including the cornu ammonis (CA) and dentate gyrus (DG) subregions, is a brain area highly susceptible to seizure-like activity (SLA). Most studies conducted in vivo have been performed in a single hippocampal subregion. In our study, we used the high [K^+^] (HK^+^) model of SLA to investigate the role of oscillatory activity in predicting SLA and in its modulation by anti-epileptic drugs in the three hippocampal subregions (CA1, CA3 and DG). For this, we recorded spontaneous local field potentials (LFPs) in CA1, CA3 and DG subregions from mouse hippocampal slices. We find that the oscillatory activity in the 20 second pre-ictal period is significantly different compared to the oscillatory activity in the absence of SLA or to a more distant period from the ictal event. A classification algorithm revealed that the oscillatory dynamics, particularly in the CA1 subregion, can predict the emergence of an ictal event with high accuracy. Furthermore, oscillatory activity is differentially modulated by anti-epileptic drugs in the different hippocampal subregions. We found that diazepam and carbamazepine modulated the oscillatory activity significantly greater in the CA3 and DG subregions, compared to CA1. Imaging of neuronal activation in the *ex vivo* model of seizure-like activity, using the Fos protein as an activity marker, revealed a similar subregion-dependent differential modulation following diazepam and carbamazepine perfusion. Therefore, while oscillatory activity in the pre-ictal period in the CA1 subregion can better predict the emergence of ictal events, anti-convulsant drugs have a stronger effect in oscillatory activity in the CA3 and DG subregions.

## Introduction

Epilepsy is a very common and serious neurological disorder, characterized by the occurrence of spontaneous recurrent seizure events. Seizure activity is a temporary manifestation of abnormally synchronized and excessive neuronal firing in the brain [1]. The etiology of seizure events is not well understood but associated with an imbalance of excitation-inhibition in neuronal networks [2,3].

Synchronous oscillatory activity is a form of correlated neuronal firing important for proper performance in cognitive tasks [4]; however, hypersynchrony is observed in both in *in-vivo* rodent models of epilepsy and in epileptic patients [5,6]. In rodents, epileptic activity exhibits increased power in low-(delta (δ) and theta (θ)) and high-frequency (gamma (γ)) oscillations in the hippocampus (HPC) [7–9]. High-frequency oscillations have also been observed in *ex vivo* models of seizure-like activity (SLA) [10]. These alterations in oscillatory activity are thought to contribute to the cognitive and behavioral deficits associated with epilepsy [11].

Considerable research efforts currently aim at identifying electrophysiological biomarkers that can be used to predict the emergence of seizures [12–17]. Seizure prediction models have focused on distinguishing between interictal and pre-ictal activity based on either the duration of each period or some time-frequency features of EEG signals [18]. Predictive models in animal models of epileptiform activity, either *in vivo* or *ex vivo*, are rare [19,20] and render the detailed study for predictive capabilities of various features difficult and fragmented.

Current pharmacotherapy targets mechanisms that affect the excitation and inhibition balance in the brain. Examples of anti-seizure mediations include diazepam, a benzodiazepine, and carbamazepine, a sodium channel blocker [21]. It is well established that both diazepam and carbamazepine reduce epileptiform discharges, both *in vivo* and in brain slices [22,23]. These two different drugs differentially affect the neuronal oscillations. Diazepam reduces all frequency bands while carbamazepine is more effective in the high frequency oscillations [24–26].

Finally, a few studies have identified a differential susceptibility and adaptations among the CA1 subfields and DG [27–29]. However, most studies have primarily focused on investigating one subregion at a time, therefore, very few data exist comparing the properties of epileptiform activity and the response to anti-epileptic drugs across subregions.

In this study, we aim, on one hand, to investigate whether pre-ictal oscillatory activity can predict the emergence of ictal events in an *ex vivo* model of seizure-like activity, and, on the other hand, to determine possible differences among the subregions of hippocampal formation in predicting ictal events. Furthermore, we investigated the effect of two well-known anti-epileptic drugs on oscillatory activity in all three subregions. For this, we use a well-established *ex vivo* model of SLA which is produced by augmenting the concentration of extracellular potassium ions [K^+^] in the perfusing solution bathing the hippocampal slice, promoting the development of both interictal and ictal-like activity patterns [30,31]. Our results show differential capabilities for predicting ictal events and for the effects of anti-epileptic drugs on oscillatory activity in the CA1, CA3 and DG regions.

## Materials and Methods

### Animals and Housing

Male and female C57Bl/6 mice, 30–60 days of age, were used for all experiments. Mice were group-housed (3 to 4 per cage) and provided with food (standard mouse chow) and water *ad libitum*. The vivarium rooms were under a 12 h light/dark cycle (light on at 7:00 am) with controlled temperature (21 ±°C). All animal procedures comply with the ARRIVE guidelines and were performed according to the European and National ethical standards. All experiments were approved by the Department of Biology, University of Crete-Experimental Protocol Committee.

### Brain Slice Preparation

Mice were euthanized under halothane anesthesia. The brain was quickly removed and placed in ice cold oxygenated artificial cerebrospinal fluid (aCSF) (95%O_2_/5% CO_2_) containing (in mM): 125 NaCl, 3.5 KCl, 26 NaHCO_3_, 1 MgCl_2_ and 10 glucose (pH =7.4, 310 mOsm/l). After the brain was blocked, the part containing the hippocampus was glued onto the stage of a vibratome (Leica, VT1000S, Leica Biosystems GmbH, Wetzlar, Germany). Coronal brain slices (400μm) containing the dorsal hippocampus (bregma -1.34 to -2.06mm) were placed in a submerged chamber containing oxygenated (95% O_2_/5% CO_2_) aCSF (in mM): 125 NaCl, 3.5 KCl, 26 NaHCO_3_, 2CaCl_2_, 1 MgCl_2_ and 10 glucose (pH = 7.4, 315 mOsm/l) at 36,6 °C. The brain slices were allowed to recover for at least 1 h in this chamber before recordings began.

### Local Field Potential (LFP) Recordings and Data Analyses

The slices were placed in a temperature-controlled slice chamber at 32^°^C with 95% O_2_-5% CO_2_ under a stereoscope continuously perfused initially with control aCSF containing (in mM): 125 NaCl, 3.5 KCl, 26 NaHCO_3_, 2CaCl_2_, 1 MgCl_2_ and 10 glucose (pH = 7.4, 310 mOsm/l), followed by HK^+^ aCSF (aCSF containing, in mM: 125 NaCl, 7.5 KCl, 26 NaHCO_3_, 1 MgCl_2_, 2 CaCl_2_ and 10 glucose (pH = 7.4, 310 mOsm/l). To evaluate the effect of known anti-epileptic drugs, diazepam (a GABA-A receptor agonist, C_DZP_= 2μM) or carbamazepine (a sodium channel blocker, C_CBZ_= 100μM) was diluted in HK^+^ aCSF. Diazepam was acquired from the Pharmacy of the University General Hospital in Heraklion as a 5 mg/ml solution. CBZ was purchased from Sigma-Aldrich.

The recording electrode (1-2MΩ resistance) was filled with 3M NaCl and was positioned in CA1, CA3 or DG regions of distinct hippocampal slices. Spontaneous LFPs were amplified using the EXT-02F amplifier (National Instruments), digitized with ITC-18 (Instrutech, Inc.) and recorded in a computer running Windows10 with WinWCP software (Stratchclyde electrophysiology software). In order to study the oscillatory activity, the selectable high pass filter was set at 0 Hz, while the signal was low-pass filtered at 300Hz.

To analyze the oscillatory activity, the time series of the voltage traces was transformed into frequency domains using a fast Fourier Transform (FFT) algorithm. The Power Spectral Density (PSD) was estimated for each frequency domain in logarithmic scale. The following frequency domains were analyzed: Delta (δ): 1–4 Hz, Theta (θ): 4–7 Hz, Alpha (α): 8–12 Hz, Beta (β): 13–30 Hz, Gamma (γ): 30–80 Hz and High Gamma (hγ): 80–200 Hz. All analysis was generated using custom-written procedures in MATLAB-R2023 (The MathWorks, Inc.).

### Detection of ictal events

The beginning of an ictal event was defined when the voltage signal was higher than 5σ above the average of the voltage signal (Pfammatter et al., 2018). An event ended when the signal decreased 4σ below the mean of the voltage signal. Seizure durations were manually measured as periods in which this rise crossed and remained above the threshold of 5σ for longer than 2sec. We employed artifact rejection techniques to remove identified signals that were 15-fold larger in amplitude than the average spike amplitude (Sorokin et al., 2017).

### Fos Immunochemistry Protocol

Hippocampal slices (250μm), prepared as described above, were placed in a submerged chamber containing oxygenated control aCSF and were allowed to equilibrate for 1 hr. Next, some of the slices were placed in a submerged chamber containing oxygenated HK^+^ aCSF and some other slices were placed in a submerged chamber containing oxygenated HK^+^ aCSF plus diazepam (C_DZP_= 2μM) or carbamazepine (C_CBZ_= 100 μM) for 25 min. Following this incubation, all slices were transferred to different chambers containing oxygenated control aCSF for another 1h. Finally, the slices were fixed in 4% paraformaldehyde solution (in different chambers) for 1h and, subsequently, washed with 0.1% PBS-Tween (PBS-T) and blocked overnight with 4% fetal bovine serum and 0.1% Triton in PBS-T. The next day, the slices were incubated with a primary antibody (c-Fos Antibody (E-8):sc-166940, Santa Cruz Biotechnology, Inc., 1:50 dilution) in 4% FBS and 0.1% Triton in PBS-T for 48 hours at 4°C. Afterwards, the slices were exposed to a secondary antibody (Goat Anti-Mouse IgG (H + L), CF®568, Biotium), were stirred for 5h at RT, mounted onto slides, and coverslipped with Mounting Medium with DAPI (ab104139) (Abcam plc.).

### Imaging of Fos-stained slices

The slices were imaged in a custom-built laser raster-scanning, multiphoton microscope (Suppl. Fig. 1) that is using a 1030 nm fs laser (Flint, Light Conversion, Vilnius, Lithuania) as an excitation source, passing through a pair of galvanometric mirrors (6215 H, Cambridge Technology, Bedford, MA, USA) before entering into an inverted microscope (Axio Observer Z1, Carl Zeiss, Jena, Germany) [32,33]. The beam is then reflected by a short pass dichroic mirror (FF700-SDi01, Semrock, Rochester, NY, USA) placed at the turret box of the microscope and is focused into the sample with a 20x 0.8NA objective-lens (Plan-Apochromat 20x/0.8NA, Carl Zeiss) or with a 40x 1.3NA objective-lens (Plan-Apochromat 40x/1.3NA, Carl Zeiss). The emitted fluorescence is collected by the same objective and is filtered by a short pass filter (FF01-680/SP, Semrock) to ensure that no laser light is reaching the detectors. Then a band-pass filter (FF1-562/40, Semrock) followed, which allowed passing the wavelengths in the range of 562 ± 20 nm, before reaching a detector, based on a photomultiplier tube PMT-blue (H9305-04, Hamamatsu, Hizuoka, Japan). The FF1-562/40 was used to detect the 2p-F from Fos antibody. Coordinate motion of the galvanometric mirrors and the detector for image acquisition was performed using custom-built Labview (National Instruments, Austin Texas, USA) software. The collage images were created by coordinate motion of the motorized sample stage of the microscope and image acquisition, using the Labview software.

### Automated Counting of Activated Cells

All images were processed using the open-source ImageJ software, with the Fiji distribution (National Institute of Health and University of Wisconsin, USA) before used for cell counting. CellProfiler 4.2.5 (http://www.cellprofiler.org/index.shtmL) was used for automated counting the Fos expressing cells. All images were of the same total area covered by cells of 1200μm^2^ for each subfield. The CellProfiler software was first used to measure the range of a typical diameter of cells 15-140, in pixel units. The Threshold method used was Minimum Cross-Entropy with lower and upper bounds 0.15-1.0, smoothing scale 1.3488 and correction factor 1.0 (Suppl. Fig.2).

### Statistical Analysis

Repeated measures or one-way ANOVA analysis were performed as appropriate, followed by the post hoc Tukey test. In cases when only two groups were tested (Fig. 1) t-test was used. Statistical analysis was performed with GraphPad Prism.

**Figure 1.**
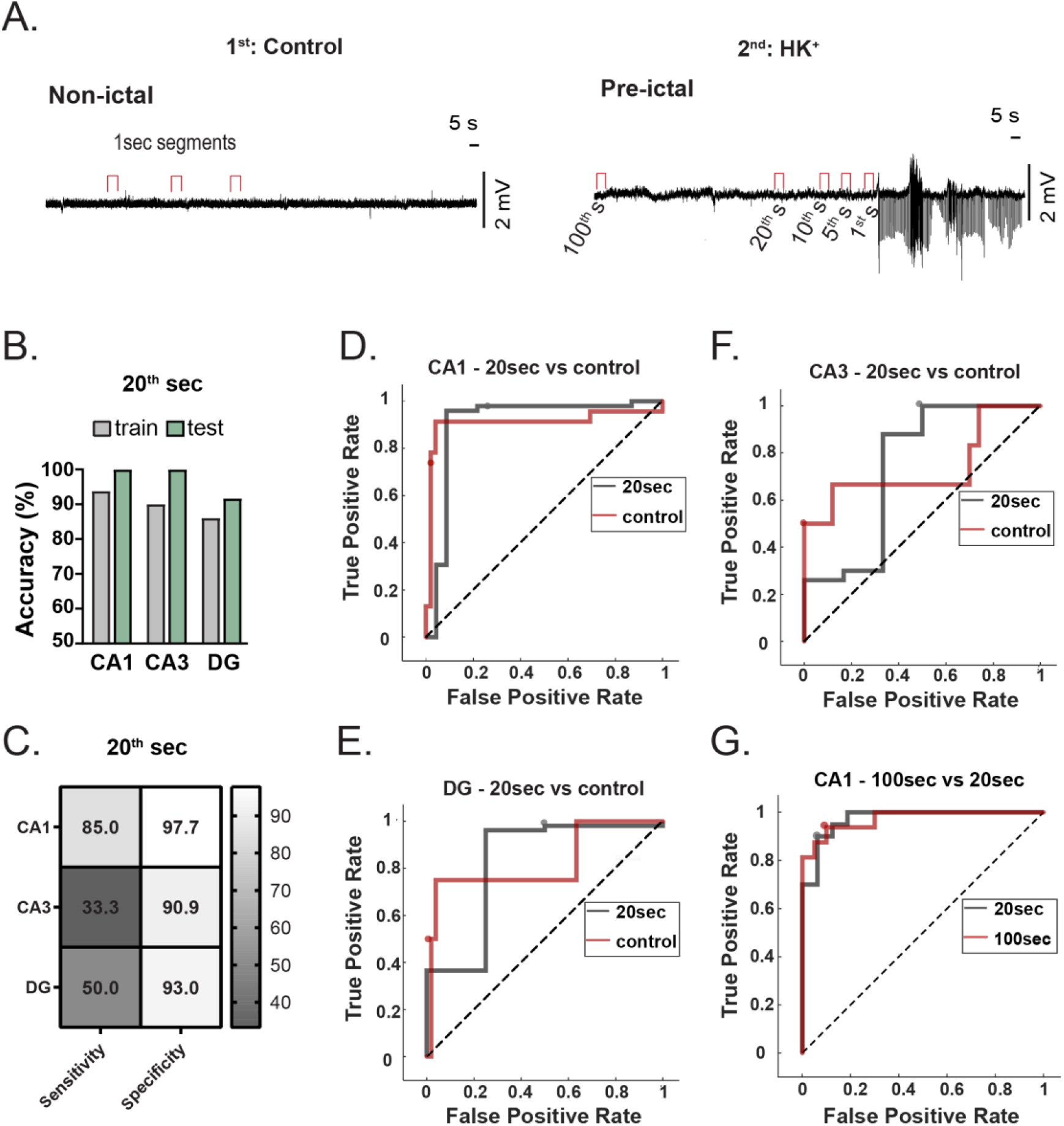
Prediction of SLA based on the spectral power of oscillatory activity during the 20^th^ second of pre-ictal activity. (A) Representative traces of the voltage signal under control (non-ictal) conditions (left) and under HK aCSF (right). The presence of SLA is evident under HK aCSF. The 100^th^, 20^th^, 10^th^, 5^th^ and 1^st^ second of the pre-ictal condition that are analyzed are indicated. (B) Total accuracy following training and testing of Linear Discriminant models using the values from the spectral power of oscillatory activity in the 20^th^ second before an ictal event and respective seconds under non-ictal conditions for the CA1, CA3 and DG regions. (C) Sensitivity and specificity of the linear discriminant models for the CA1, CA3 and DG regions. (D-F) ROC curves for the linear discriminant models in CA1 (C), CA3 (D) and DG (E) regions. (G) ROC curve for the linear discriminant model in CA1 region using the values from the spectral power of oscillatory activity in the 20^th^ second before an ictal event and the 100^th^ second before the ictal event.

## Results

### Pre-ictal analysis of the oscillatory activity in the CA1, CA3 and DG regions

Perfusion of HK aCSF resulted in induction of SLA in all three hippocampal subregions (Suppl. Fig. 3). Following identification of an ictal discharge, we analyzed the spectral power of different frequencies (δ, θ, α, β, γ and hγ) from 1sec segments at distinct time points relative to the start of ictal discharges. Specifically, we analyzed the 20^th^, 10^th^, 5^th^ and 1^st^ second prior to the start of the ictal discharge (pre-ictal), as well as various 1sec segments of spontaneous activity in the control aCSF condition (non-ictal) in the same brain slice (Tables 1, 2, 3). The power of most frequencies was significantly increased in the 20^th^ and 10^th^ second before the ictal event, compared to control spontaneous activity in the CA1 (Table 1). In the CA3 region, power in the θ, α, β, γ and hγ domains was increased between the 5^th^ second (primarily) before the ictal event and the control spontaneous activity (non-ictal) (Table 2). In the DG region, a similar increase in power was observed in the θ, α, β, and hγ domains during the 5^th^ second (primarily) before the ictal event when compared to the control spontaneous activity (non-ictal) (Table 3). Thus, it seems that modulation of oscillatory activity is more substantial in the CA1 region, compared to CA3 and DG regions. Given that the 20^th^ second was the more distant time point which exhibited differences in the power of oscillations compared to the control, non-ictal, condition, we proceeded with this time point for the prediction analysis of the ictal events below.

**Table 1.** Temporal analysis of the power of oscillations across all frequency bands between a non-ictal-segment and the pre-ictal 1-second segments in CA1.

|  | 20<sup>th</sup> | 10<sup>th</sup> | 5<sup>th</sup> | 1<sup>st</sup> | Non-ictal | One-way ANOVA |
| --- | --- | --- | --- | --- | --- | --- |
| $\delta$ | 3.17E+05** | 3.77E+05** | 2.69E+05 | 2.87E+05* | 1.64E+05 | $F_{(4, 123)}=4.66$,<br>$p^{**}=0.0015$ |
| $\theta$ | 1.79E+05** | 2.09E+05** | 1.94E+05** | 1.66E+05** | 7.81E+04 | $F_{(4, 123)}=6.79$,<br>$p^{****}<0.0001$ |
| $\alpha$ | 1.54E+05*** | 1.64E+05* | 1.62E+05** | 1.62E+05*** | 7.39E+04 | $F_{(4, 123)}=7.88$,<br>$p^{****}<0.0001$ |
| $\beta$ | 4.77E+05**** | 4.34E+05* | 4.62E+05** | 4.82E+05**** | 1.36E+05 | $F_{(4, 123)}=9.34$,<br>$p^{****}<0.0001$ |
| $\gamma$ | 1.11E+06 | 2.19E+06**** | 1.95E+06**** | 1.93E+06**** | 8.84E+05 | $F_{(4, 123)}=15.41$,<br>$p^{****}<0.0001$ |
| $h\gamma$ | 1.96E+06**** | 1.68E+06**** | 1.78E+06**** | 1.71E+06**** | 5.63E+05 | $F_{(4, 123)}=22.63$,<br>$p^{****}<0.0001$ |

**Table 2.** Temporal analysis of the power of oscillations across all frequency bands between a non-ictal-segment and the pre-ictal 1-second segments in CA3.

|  | 20<sup>th</sup> | 10<sup>th</sup> | 5<sup>th</sup> | 1<sup>st</sup> | Non-ictal | One-way ANOVA |
| --- | --- | --- | --- | --- | --- | --- |
| $\delta$ | 3.69E+05 | 1.73E+05 | 3.27E+05 | 2.34E+05 | 2.26E+05 | $(F_{(4, 99)}=1.23$,<br>$p=0.39$ |
| $\theta$ | 1.87E+05 | 1.78E+05 | 2.24E+05 *** | 1.42E+05 * | 9.95E+04 | $F_{(4, 99)}=5.449$,<br>$p^{***}<0.0005$ |
| $\alpha$ | 1.61E+05 | 1.43E+05 | 1.78E+05 ** | 1.44E+05 | 9.51E+04 | $F_{(4, 99)}=4.58$,<br>$p^{**}<0.0019$ |
| $\beta$ | 5.12E+05** | 3.79E+05 | 4.64E+05** | 3.69E+05*** | 2.38E+05 | $F_{(4, 99)}=7.92$,<br>$p^{****}<0.0001$ |
| $\gamma$ | 1.29E+06 | 1.48E+06* | 1.33E+06* | 1.27E+06* | 2.03E+06 | $(F_{(4, 99)}=5.14$,<br>$p^{**}<0.0005$ |
| $h\gamma$ | 2.90E+06* | 2.83E+06*** | 3.28E+06**** | 3.17E+06*** | 1.67E+06 | $F_{(4, 99)}=10.5$,<br>$p^{****}<0.0001$ |

**Table 3.** Temporal analysis of the power of oscillations across all frequency bands between a non-ictal-segment and the pre-ictal 1-second segments in DG.

|  | 20<sup>th</sup> | 10<sup>th</sup> | 5<sup>th</sup> | 1<sup>st</sup> | Non-ictal | One-way ANOVA |
| --- | --- | --- | --- | --- | --- | --- |
| $\delta$ | 3.95E+05* | 2.25E+05 | 3.35E+05* | 2.96E+05 | 2.02E+05 | $(F_{(4, 87)}=4.08, p^{**}=0.004)$ |
| $\theta$ | 2.07E+05*** | 1.51E+05 | 1.69E+05*** | 1.67E+05*** | 9.56E+04 | $F_{(4, 87)}=10.41, p^{****}<0.0001$ |
| $\alpha$ | 1.41E+05 | 1.25E+05 | 1.47E+05** | 1.65E+05*** | 8.85E+04 | $F_{(4, 87)}=7.18, p^{****}<0.0001$ |
| $\beta$ | 3.81E+05 | 3.71E+05 | 4.58E+05 | 5.16E+05*** | 2.30E+05**** | $F_{(4, 87)}=10.90, p^{****}<0.0001$ |
| $\gamma$ | 1.27E+06 | 1.81E+06 | 2.29E+06 | 1.69E+06 | 1.50E+06 | $F_{(4, 87)}=1.49, p=0.21$ |
| hγ | 1.77E+06 | 1.93E+06 | 2.31E+06* | 2.02E+06**** | 1.12E+06** | $(F_{(4, 87)}=8.25, p^{****}<0.0001)$ |

### Ictal event prediction based on the power of oscillatory activity during the 20sec pre-ictal period

Based on the observed differences in the power of oscillatory activity between the pre-ictal and non-ictal periods, we tested whether the spectral power of oscillatory frequencies in the pre-ictal period can predict the emergence of an ictal event. To achieve this, the spectrum power of the different frequencies at the 20^th^ second before the emergence of an ictal event (i.e. SLA), as well as in 1-sec segments under control non-ictal conditions) were used to train and test a classification algorithm using Linear Discriminant Analysis (LDA). The results indicated that SLA in CA1 region can be predicted with better accuracy, compared to SLA in the CA3 and DG regions (Fig. 1A). The sensitivity and specificity of the model were also notably better (97.7%) for the CA1 compared to that of the CA3 and DG regions (Fig. 1B-E). We also tested whether the LDA model could distinguish the power of oscillations between the 20^th^ second before SLA and a more distant 100^th^ second. We found that the model demonstrated high accuracy in the training set (81.3%), which was maintained in the test set, achieving an accuracy of 77.3%. Additionally, the model exhibited very good specificity and sensitivity (87.5% and 78% and Fig. 1F). Finally, shuffling of the data that were used to train and test the CA1 LDA model resulted in significantly poorer performance (71% train accuracy, 64% test accuracy, 33% sensitivity and 73% specificity). Therefore, our results suggest that an LDA algorithm trained on the spectral power of oscillatory activity can predict the emergence of ictal events 20 seconds before their onset, even under ictogenic conditions. This prediction is much more accurate in the CA1 region recordings.

### Effects of diazepam and carbamazepine administration on oscillatory activity

Given the importance of oscillatory activity in predicting SLA, we investigated the modulation of oscillatory activity following perfusion of diazepam or carbamazepine, two well-known anti-epileptic drugs. Application of diazepam and carbamazepine reduced SLA in the CA1, CA3 and DG regions (Suppl. Fig. 4). Upon analysis of the oscillatory activity, we found that diazepam did not have any effect in the oscillatory activity in the CA1 region (Fig. 2A), however, it did reduce the spectral power of θ, α, βand hγ frequencies in the CA3 region (Fig. 2B) and the spectral power of θ, α, β, γ, and hγ frequencies in the DG region (Fig. 2C). Similarly, carbamazepine did not have any effect in the oscillatory activity in the CA1 region (Fig. 3A), however, it did reduce the spectral power of all frequencies in both CA3 (Fig. 3B) and DG regions (Fig. 3C). Thus, our results indicate that the CA3 and DG subregions are more sensitive to modulation of oscillatory activity, following perfusion diazepam and carbamazepine.

**Figure 2.**
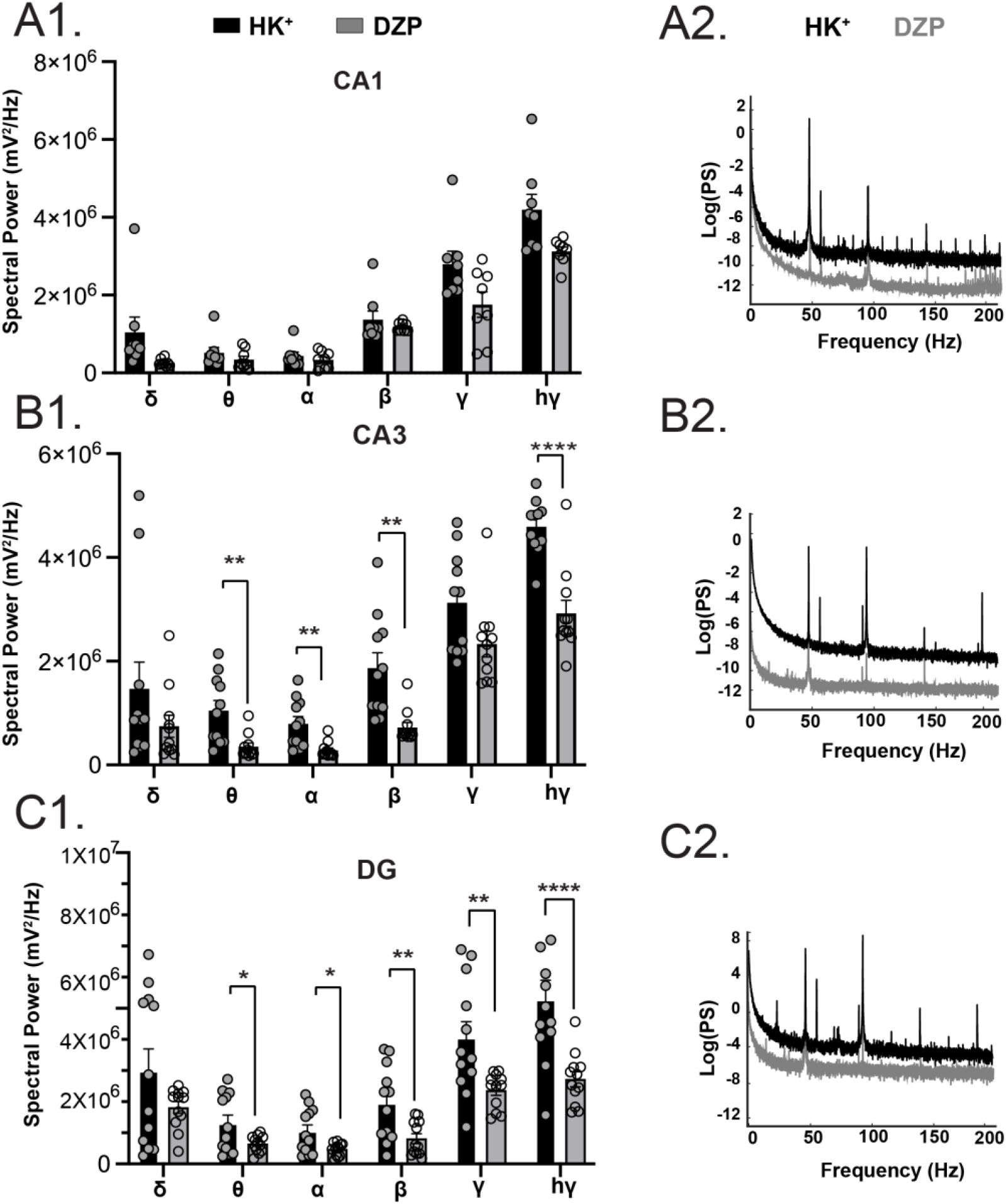
Effect of diazepam (DZP) on the oscillatory activity of the CA1, CA3 and DG subregions. (A) Bar graph (A1) and representative traces (A2) showing no effects of diazepam on the oscillatory activity in the CA1 subregion (repeated measures ANOVA, F_(1,14)_=4.59; p^*^=0.05). Post-hoc comparisons did not reveal any significant differences between HK^+^ and diazepam groups in any of the frequencies. (B) Bar graph (B1) and representative traces (B2) showing significant effects of diazepam on θ, α, β and hγ frequencies in the CA3 subregion (repeated measures ANOVA analysis, F_(1,20)_=15.74, p^**=^0.0008). Tukey’s multiple comparisons revealed a significant effect in θ (p^**^=0.005), α (p^**^=0.004), β (p^**^=0.003) and hγ (p^****^<0.0001). (C) Bar graph (C1) and representative traces (C2) showing significant effects of diazepam on the θ, α, β, γ and hγ frequencies in the DG subregion (repeated measures ANOVA analysis, F_(1,22)_=11.13, p^**^=0.003, Tukey’s multiple comparisons revealed a significant effect in the in θ (p^*^=0.03), α (p^*^=0.02), β (p^**^=0.009), γ (p^**^=0.009) and hγ (p^**^<0.002).

**Figure 3.**
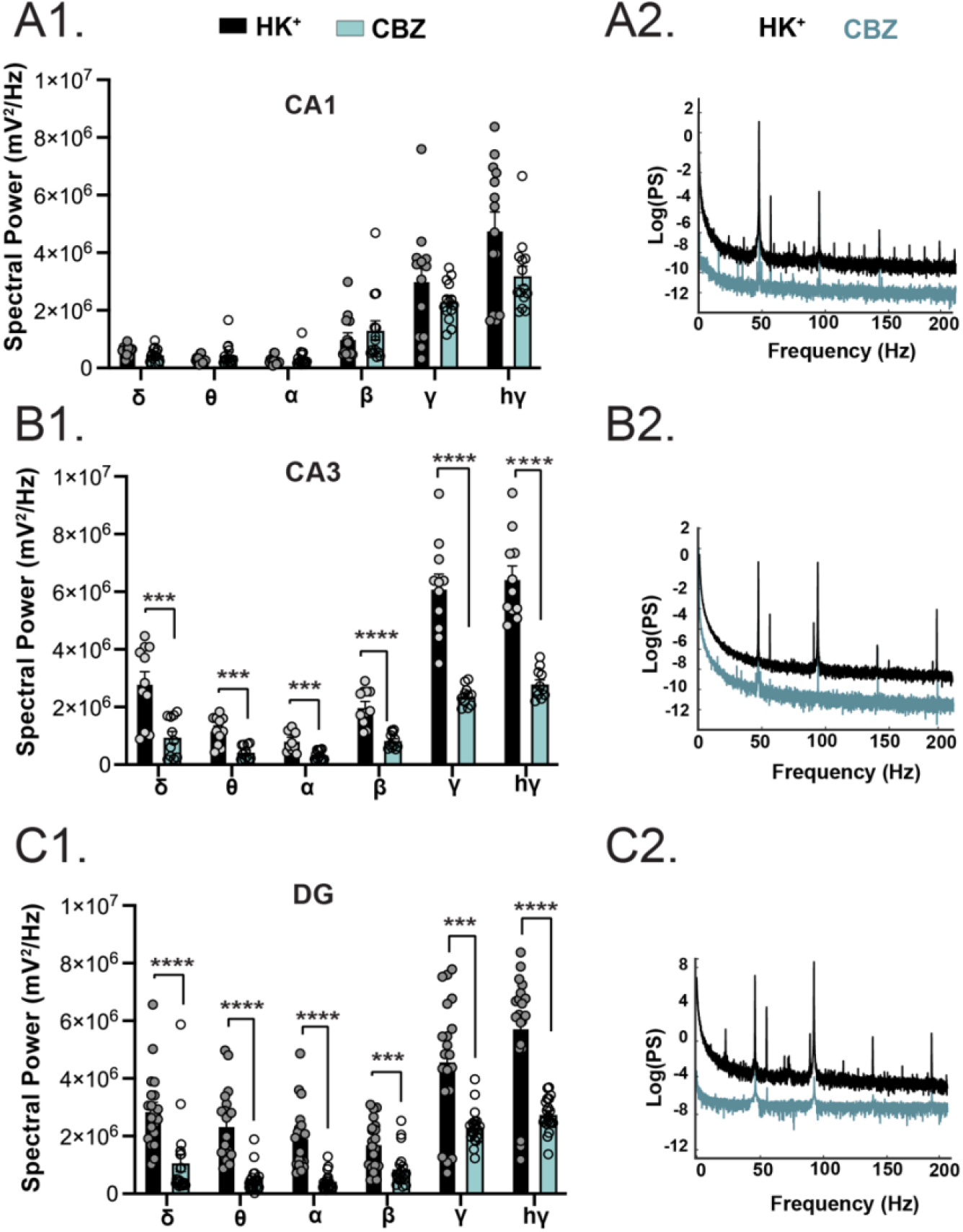
Effect of carbamazepine (CBZ) on the oscillatory activity of the CA1, CA3 and DG subregions. (A) Bar graph (A1) and representative traces (A2) showing no effects of carbamazepine on the oscillatory activity in the CA1 subregion (repeated measures ANOVA, F_(1,26)_=2.57, p=0.12) (B) Bar graph (B1) and representative traces (B2) showing significant effects of carbamazepine on the δ, β, γ and hγ frequencies in the CA3 subregion (repeated measures ANOVA analysis, F_(1,20)_=62.42, p^****^<0.0001, Tukey’s multiple comparisons revealed a significant effect in δ (p^***^=0.0009), θ (p^***^=0.0003), α (p^***^=0.0006), β (p^****^<0.0001), γ (p^****^<0.0001) and hγ (p^****^<0.0001). (C) Bar graph (C1) and representative traces (C2) showing significant effects of carbamazepine on the δ, θ, α, γ and hγ frequencies in the DG subregion (repeated measures ANOVA analysis, F_(1,38)_=96.24, p^****^<0.0001, Tukey’s multiple comparisons revealed a significant effect in δ (p^****^<0.0001), θ (p^****^<0.0001), α (p^****^<0.0001), β (p^***^=0.0007), γ (p^***^=0.0002) and hγ (p^****^<0.0001).

### Effects of diazepam or carbamazepine administration on the number of Fos expressing cells in the hippocampus using multiphoton imaging

Since the electrophysiological recordings in the different regions of the hippocampal formation were not performed concurrently, we utilized an imaging approach to further investigate differences between the CA1, CA3 and the DG. We performed Fos immunofluorescence in hippocampal slices after the administration of the same perfusing conditions used in the electrophysiological recordings. Perfusion of diazepam in the presence of HK^+^ aCSF resulted in significant reduction of the number of Fos positive cells in the CA1, CA3 and DG regions. Perfusion of carbamazepine significantly reduced the number of Fos positive cells in CA3 and DG, but not in CA1 (Fig. 4).

**Figure 4.**
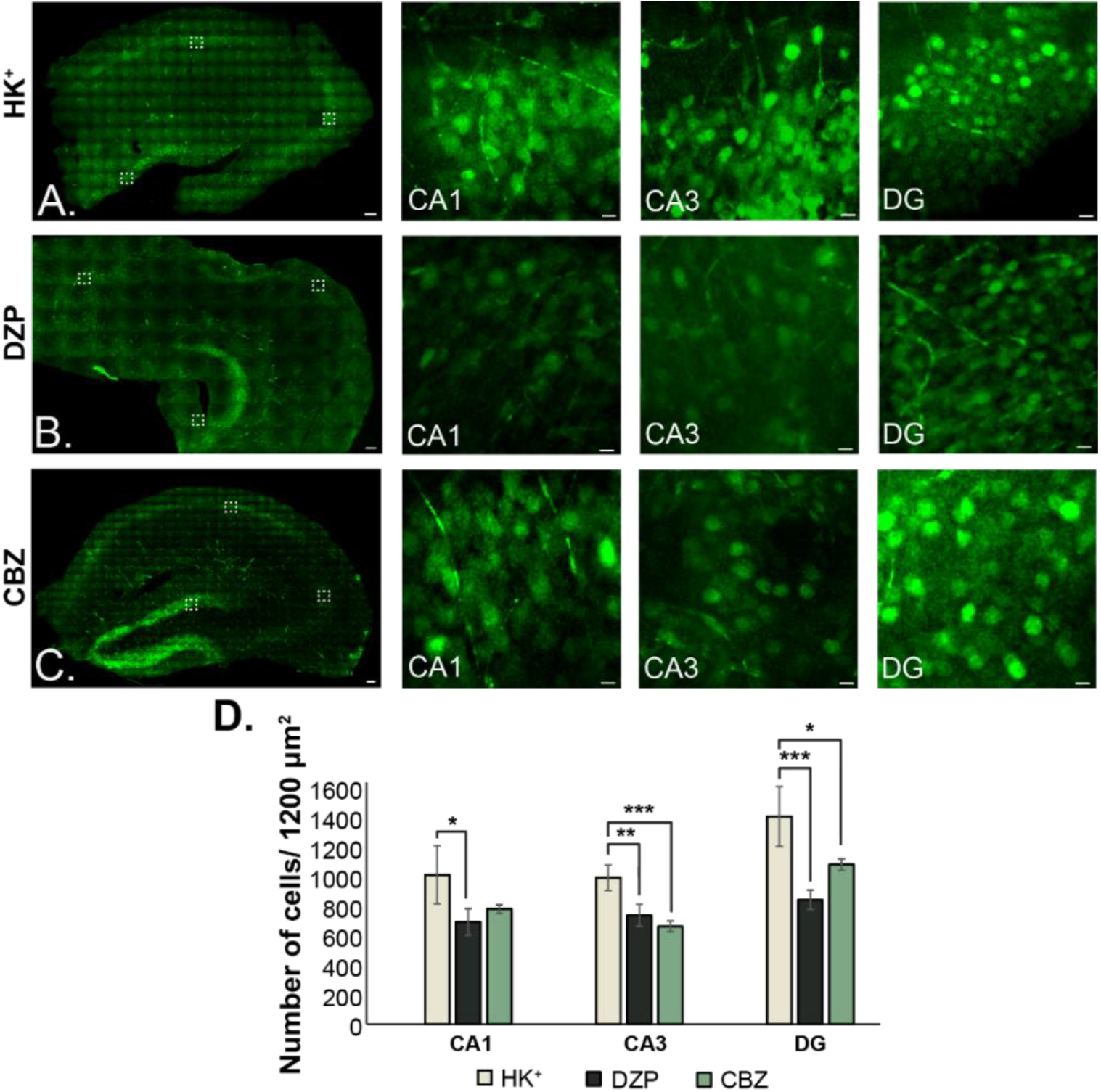
Expression of c-Fos in the HPC. (A-C) Representative multiphoton images after the perfusion of (A) HK^+^, (B) diazepam (DZP) and (C) carbamazepine (CBZ) aCSF (scale bar 100μm for the full one hemisphere hippocampus and 10μm for CA1, CA3 and DG separately). Dotted squares display representative images for the 3 regions. (D) Number of activated cells that were identified after the different perfusing conditions in the 3 subfields. (CA1: one way ANOVA (F_(2,16_)=5.53, p^**^=0.014; post-hoc Tukey test DZP<HK^+^ p^*^=0.02, CBZ<HK^+^ p=0.14; CA3: one way ANOVA (F_(2,16)_=6.73, p^***^=0.007; post-hoc Tukey test DZP<HK^+^ p^**^=0.01, CBZ<HK^+^ p^***^=0.001; DG: one way ANOVA (F_(2,16)_=7.77, p^***^=0.004; ; post-hoc Tukey test DZP<HK^+^ p^***^=0.001, CBZ<HK^+^ p^*^=0.04) (n=6 slices per group).

## Discussion

Our study revealed significant differences across the CA1, CA3 and DG subregions of neuronal oscillations in both predicting the emergence of seizure-like activity and its modulation by anti-epileptic drugs. Firstly, our results showed that the oscillatory activity in the 20-sec pre-ictal period are significantly different compared to the oscillatory activity under non-ictogenic conditions or at a more distant time before the ictal event. Additionally, the oscillatory activity during the 20^th^ second could predict the emergence of SLA with high accuracy and excellent sensitivity/specificity, in the CA1 region. Secondly, anti-epileptic drugs (diazepam and carbamazepine) modulated both low and high-frequency neuronal oscillations, primarily in the CA3 and the DG.

### Ictal event prediction

In this study, we highlight the significant differences in oscillatory dynamics within the pre-ictal period, as well as across the pre-ictal and non-ictal periods. Therefore, based on these differences, we investigated whether we could predict the emergence of ictal events based on the power of different oscillations and found that prediction is possible with high accuracy, specificity and sensitivity, particularly in the CA1 region.

Most predictive models have utilized clinical datasets obtained from either intracranial EEG or scalp EEG and aim to distinguish between the interictal and pre-ictal periods [14] or between the early and late pre-ictal states [12]. Many human studies utilize metrics related to the frequencies of oscillatory dynamics based on EEG recordings with duration 3min – 60 min before seizure onset in order to prospectively predict seizure events [16,17,34,35]. However, oscillatory activity information extracted from shorter duration intervals could be advantageous and be programmed in a continuous on-line analysis manner in future predictive algorithms. In animal models, there are only a couple of studies applying predictive algorithms for epileptic discharge emergence. The studies have used various statistical metrices of spiking activity [19] or ion concentration changes [20] to predict the emergence of an epileptic event. In our study, we wanted to determine whether oscillatory activity, which is primarily used in human studies, in 1sec duration intervals could be used to predict the emergence of an ictal event in our ex vivo model of SLA. We have found that, during the 20^th^ second before an ictal event in the CA1, oscillatory activity in 1sec intervals could predict the imminent occurrence of an ictal event with high accuracy and excellent sensitivity/specificity. Conversely, prediction accuracy based on the oscillatory dynamics in the CA3 and DG subregions was below 90% and the sensitivity/specificity quite poor. Consequently, our results suggests that the oscillatory dynamics could predict the emergence of an ictal event in a region-specific manner. Further experiments investigating these types of predictive model in *in vivo* animal models of epilepsy could provide valuable insights for developing further patient-specific analyses.

### Differential modulation of oscillatory dynamics by diazepam and carbamazepine

Perfusion of the HK^+^ aCSF elicited SLA in all three subregions of the hippocampal formation, as expected [36,37]. Both slow and fast neuronal oscillation were significantly enhanced under ictogenic conditions (i.e. in the presence of HK^+^ aCSF). These results align with previous reports that observed that the peak rates of the mid and high frequency domains exhibit higher levels during ictogenic conditions [38–40]. Both diazepam and carbamazepine reduced the oscillatory activity in the high frequencies, however, only carbamazepine reduced the oscillatory activity in the low frequency delta. Similar to our study, diazepam has been found to reduce theta and gamma frequencies [41]. *In vivo* studies in wild-type animals have found that diazepam enhances slow oscillations [25,42]. In our study, we did not identify any changes in the slow (delta) oscillations, suggesting that the in vivo reported changes are likely due to effects of diazepam on long-range connectivity. Carbamazepine is known to reduce mid and high-frequency oscillations [24,43], however, we did not find any studies addressing the effect of carbamazepine in slow oscillations in the hippocampal formation.

### Spatial differences in the hippocampal formation

In our study, we have identified several differences among the 3 regions of the hippocampal formation: CA1, CA3 and DG. First, we identify that the pre-ictal oscillatory dynamics can predict the emergence of an ictal event with much better accuracy, sensitivity and specificity in the CA1 region, compared to CA3 and DG regions. Second, the effects of diazepam and carbamazepine in spontaneous spiking activity is not significant in the DG region. Third, the modulation of oscillatory dynamics by diazepam and carbamazepine is much greater in CA3 and DG region, while not significant in the CA1 region. The regional differences could depend on the molecular, biophysical and cellular variances that exist among the distinct local neuronal networks in CA1, CA3 and DG. For example, the cellular transcriptome and proteome among CA1, CA3 and DG regions exhibits regional variations [44,45]. Seizure induction results in changes in miRNA expression with regional variability [46]. Finally, differential sensitivity of the CA1 and CA3 regions have also been identified regarding the amount of neuronal loss in hippocampal sclerosis [47]. Since no specific mechanisms that can explain the regional variability currently exists, future studies should focus on identifying the role of putative molecular or biophysical mechanisms on the regional differences that are observed in epileptic activity modulation.

### Expression of Fos protein and oscillatory dynamics

In our study, the expression of c-Fos was significantly decreased in the CA3 and DG subfields following the perfusion of diazepam or carbamazepine, but not in CA1. Consequently, our results suggest that Fos activation could be modulated by diazepam and carbamazepine in a region-dependent manner. The adaptation pattern observed with Fos modulation (more in CA3 and DG and less in CA1) resembled the pattern observed in oscillatory dynamics modulation in the three hippocampal subregions. Correlated changes of Fos expression and oscillatory dynamics have been identified in other reports as well [48,49]. Taken together, the expression of c-Fos could possibly serve as a molecular biomarker for the network synchronicity.

### Limitations of the study

An important limitation of our study, regarding the effects of diazepam and carbamazepine, is the fact that we did not perform simultaneous recordings in the three regions investigated (CA1, CA3 and DG). However, we were able to do that in our imaging investigation of c-fos activation, although in this case due to technical reasons we had to use brain slices of different thickness compared to the thickness used in the electrophysiology experiments.

## Acknowledgements

This work was supported by the FLAG-ERA JTC 2017 project EPIGRAPH, funded nationally by GSRT (Greece, National Project Code: T18EPA2-00008) (ES) and the European Union’s Horizon 2020 research and innovation program under the Marie Skłodowska-Curie grant agreement No 101007926 (KS).

