## Supplementary material for "Region-specific modulation and predictive potential of the oscillatory dynamics in an *ex vivo* model of ictogenesis in the CA1, CA3 and dentate gyrus"

### Supplementary methods

#### Automated Counting of Activated Cells

All images were analyzed and processed using an open-source image analysis software ImageJ, before automated counting of activated cells after the different perfusing conditions using software CellProfiler 4.2.5.

Below the step-by-step analysis procedure for cell counting is described. First, the regions of interest from 2PEF images of fixed mouse brain slice were selected at which the cells covered total surface area of 1200μm$^2$. The dimensions of the excited and detected squared field-of-view of brain slice were 300μm x 300μm by a 20x objective lens and were 150μm x 150μm by a 40x objective lens. Therefore, the number of cells counted was recorded in a selected set of 4 and 8 squares by a 20x and 40x objective lens respectively. Then, an appropriate image segmentation technique should be implemented for an accurate cell counting. This technique subdivides each image into multiple sub-images, which correspond to cells, depending on the range of typical diameter and gray values of sub-images using threshold. The minimum and the maximum diameter of cells were manually measured and defined 15 and 140 in pixel units respectively. The Threshold method used was Minimum Cross-Entropy with measured lower and upper bounds on normalized pixel intensities 0.15 and 1.0 respectively. This method considers the image foreground and background as two different signal sources, above and below an intensity threshold. Here, specifically, below the 0.15 normalized pixel intensity was considered as background, while above this value was considered as foreground. The Cross-entropy thresholding is formulated as the minimization of distance of the distributions of the observed image and of the reconstructed image (Al-Ajlan & El-Zaart, 2009) and the minimum cross-entropic algorithm selects the threshold which minimizes the measure of data consistency entropy between the segmented image and the original image (Sankur, 2004). Therefore, only if the assumptions of the range of normalized gray values 0.15-1.0 and of diameter 15-140 in pixel units were valid, the resulted sub-images were identified as cells. The reconstructed image was processed using smoothing scale 1.3488 and correction factor 1.0.

### Supplementary Figures

#### Custom built, laser raster-scanning, multiphoton microscope for two-photon excited fluorescence (2p-F) microscopy

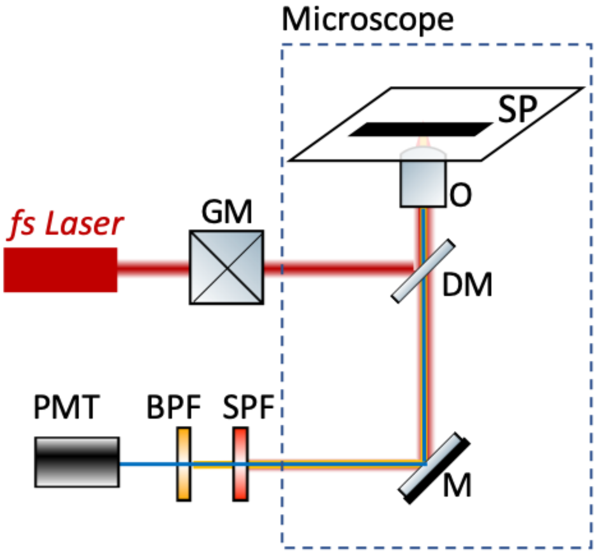


*Suppl. Figure 1* ***a) Schematic representation of the experimental setup used for 2p-F imaging microscopy of c-Fos antibody.*** *Abbreviations: GM, galvanometric mirrors; DM, dichroic mirror that reflects wavelengths longer than 700 nm and let pass wavelengths shorter than 700 nm; OL, objective lens; SP, sample plane; SPF, short-pass filter that let pass wavelengths shorter than 680 nm; BPF, band pass filter that let pass wavelengths in the range 562 ± 20 nm; PMT, photomultiplier tube.*

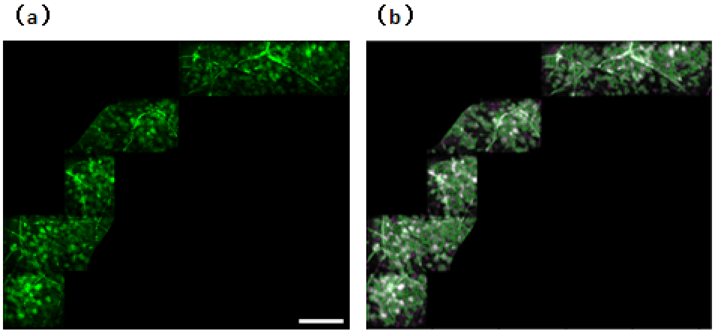


(c)

| | |
|---|---|
| # of accepted objects | 776 |
| 10th pctile diameter | 19.1 pixels |
| Median diameter | 32.5 pixels |
| 90th pctile diameter | 52.5 pixels |
| Area covered by objects | 15.00% |
| Thresholding filter size | 1 |
| Threshold | 0.15 |
| Declumping smoothing filter size | 10.1 |
| Maxima suppression size | 10.5 |

*Suppl. Figure 2* ***(a) the observed 2PEF and (b) its reconstructed image of a region of interest of a fixed mouse brain slice.*** *The 2PEF signal was collected in the backward* *detection path by a 40x objective lens. The number of cells counted was recorded in a selected set of 8 squares, covering a total surface area of 1200μm$^2$. The reconstructed image depicts the identified and discarded cells used for counting, which are indicated by green and purple circles around cells respectively. The reconstructed image was derived after an appropriate image segmentation and a thresholding technique. The range typical diameter of cells was selected 15-140 in pixel units for image segmentation. While, the Threshold method used was Minimum Cross-Entropy with lower and uper bounds 0.15-1.0, the smoothing scale 1.3488 and the correction factor 1.0 for thresholding.* ***(c) The resulted table with the recorded value of cell counting.*** *Image segmentation, thresholding and cell counting were implemented by using CellProfiler software. The scale bar is 100μm.*

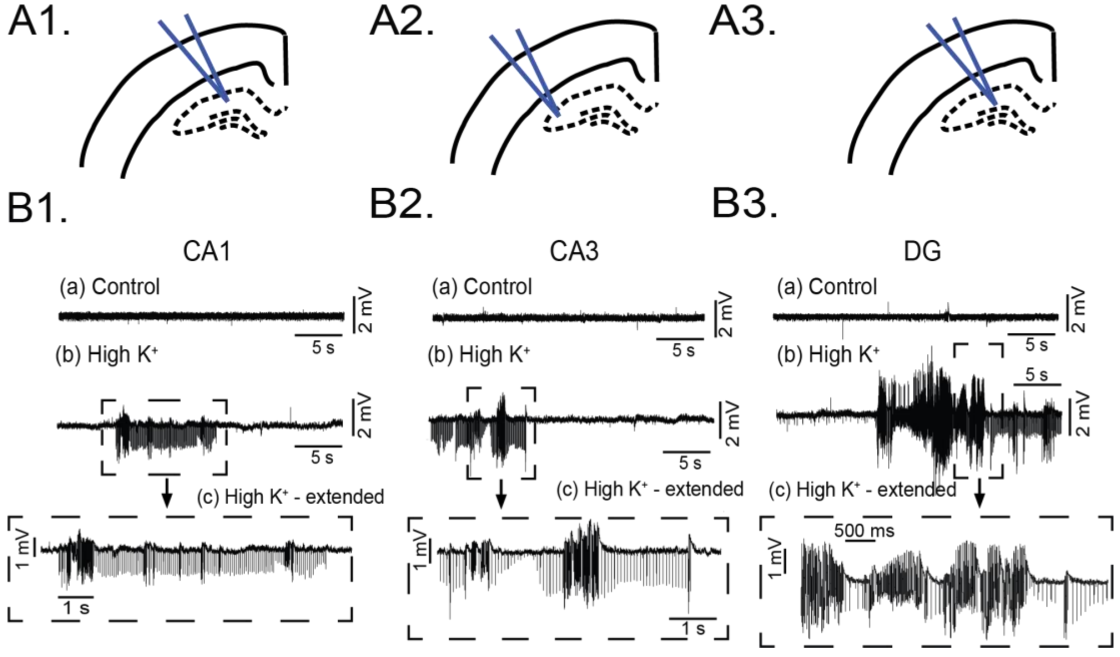


***Supplementary Figure 3. Emergence of SLA in HK$^+$ aCSF in the CA1, CA3 and DG subregions***

*(A1-3) Schematic showing the placement of electrode in the CA1, CA3 and DG subregions*

*(B1-3) Representative voltage traces following control and HK$^+$ perfusion. (B1.$_c$-B3.$_c$) Dotted lines indicate extended voltage traces showing ictal discharges.*

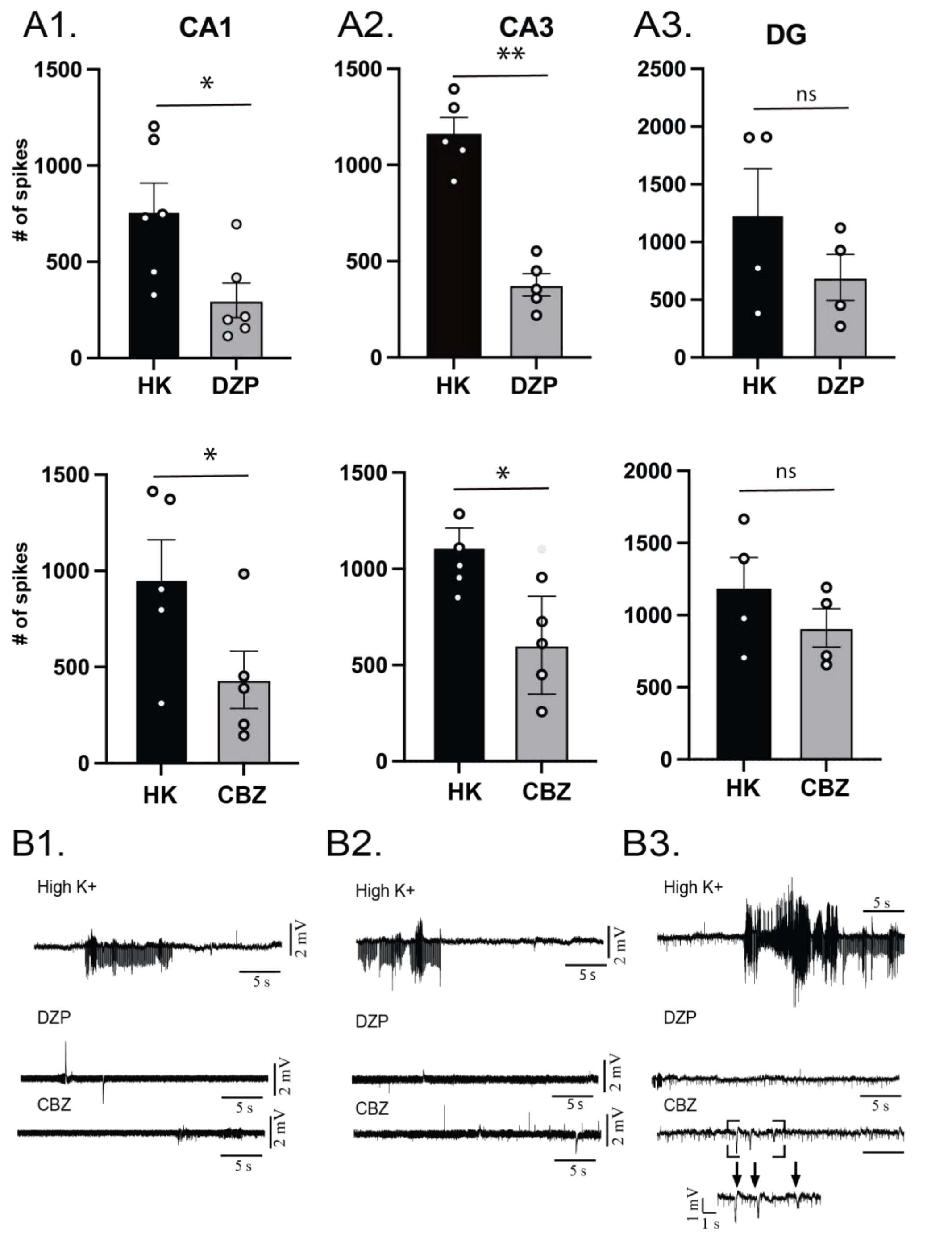


*Suppl. Fig. 4*

*A1. Effects of diazepam (DZP) and carbamazepine (CBZ) on the # of spontaneous spikes in the CA1 region. Paired t-test p=0.02 (DZP, p=0.01 (CBZ)*

*A2. Effects of diazepam (DZP) and carbamazepine (CBZ) on the # of spontaneous spikes in the CA1 region. Paired t-test p=0.002 (DZP, p=0.01 (CBZ)*

*A3. Effects of diazepam (DZP) and carbamazepine (CBZ) on the # of spontaneous spikes in the CA1 region. Paired t-test p=0.14 (DZP, p=0.35 (CBZ)*

*B1-B3. Representative traces of spontaneous activity under perfusion of HK aCSF (top), perfusion of HK + diazepam (middle) and perfusion of HK + carbamazepine (bottom). The presence of interictal spikes during perfusion of HK+diazepam and carbamazepine in the DG regions is evident (B3).*
